# tFold-Ab: Fast and Accurate Antibody Structure Prediction without Sequence Homologs

**DOI:** 10.1101/2022.11.10.515918

**Authors:** Jiaxiang Wu, Fandi Wu, Biaobin Jiang, Wei Liu, Peilin Zhao

**Author notes:** Equal contribution (Jiaxiang Wu, Fandi Wu). Work done during Fandi Wu’s internship at Tencent AI Lab.

## Abstract

Accurate prediction of antibody structures is critical in analyzing the function of antibodies, thus enabling the rational design of antibodies. However, existing antibody structure prediction methods often only formulate backbone atoms and rely on additional tools for side-chain conformation prediction. In this work, we propose a fully end-to-end architecture for simultaneous prediction of backbone and side-chain conformations, namely tFold-Ab. Pre-trained language models are adopted for fast structure prediction by avoiding the time-consuming search for sequence homologs. The model firstly predicts monomer structures of each chain, and then refines them into heavy-light chain complex structure prediction, which enables multi-level supervision for model training. Evaluation results verify the effectiveness of tFold-Ab for both antibody and nanobody structure prediction. In addition, we provide a public web service for antibody structure prediction at https://drug.ai.tencent.com/en.

## 1 Introduction

Antibodies are Y-shaped proteins produced by B cells during the immune response and play a critical role in the prevention, diagnosis, and treatment of diseases. With the development of gene sequencing technology, the cost of obtaining antibody sequences is reducing, but experimentally determining their structures is still time-consuming and expensive. Nevertheless, accurate structures are vital in analyzing the function of antibodies. Antibodies perform their functions by binding to antigens, which is mainly determined by the structure of a set of 6 loops, known as complementarity determining regions (CDRs). Accurate modeling of antibodies, especially for these CDR loops, helps researchers to better understand the antibody-antigen binding mechanism and enables the rational design of antibodies. Thus, it is urgent to develop accurate and efficient antibody structure prediction methods.

Traditional grafting-based approaches, *e.g*., ABodyBuilder [19], firstly select the template framework and then conduct CDR loop modeling and side-chain prediction. Recent methods adopt deep neural networks for antibody structure prediction. ABlooper [1] utilizes an ensemble of five E(n)-EGNNs [35] to predict the position of backbone atoms in CDR loops. DeepH3 [33] and its improved version, DeepAb [34], predict inter-residue geometric restraints with residual networks and then perform constrained energy minimization with Rosetta. NanoNet [8] firstly aligns all the training structures and then builds a 1D convolutional network to predict 3D coordinates of backbone and *C_β_* atoms. However, above methods only use neural networks to predict intermediate structures or restraints, and still rely on additional tools for full-atom structures, which limits the prediction accuracy.

In 2020, AlphaFold2 [15] achieves atomic accuracy for general protein structure prediction with a Transformer-based architecture and fully end-to-end optimization from multiple sequence alignments (MSAs) to structures. AlphaFold-Multimer [10] further extends this to structure prediction for multichain protein complexes. However, both methods heavily rely on sequence homology inputs, which can be time-consuming for large-scale sequence databases. Other methods [11, 40] turn to pre-trained language models (PLMs) to circumvent the costly MSA search, but their precision has yet to improve due to the lack of explicit sequence homologs. Yet, the performance of such end-to-end prediction methods is not extensively verified for antibody structures, especially for CDR loops and multimer sequence inputs. IgFold [31] adopts AntiBERTy [32] (pre-trained on antibody sequences) for initial features and predicts the backbone conformation in an end-to-end manner. However, side-chain structures are not explicitly considered, which can be critical for accurate prediction of CDR loops.

In this work, we propose a novel antibody structure prediction method with accurate CDR loop modeling and high inference speed. Specifically, we adopt a pre-trained language model for MSA-free prediction, and introduce a fully end-to-end architecture that firstly predicts monomer structures for heavy and light chains respectively, and then further refines them for structure prediction of heavy-light chain complexes. This allows multi-level supervision for model training with either heavy-light chain complex or single chain only. Empirical evaluation indicates that the proposed method achieves state-of-the-art performance in both antibody and nanobody structure prediction.

## 2 Related Work

### Protein Language Model

Inspired by the development of new natural language processing approach, a few protein language models (PLMs) have been developed to model individual protein sequences. PLMs learn to predict masked amino acids given their context. These models mainly use a attention-based deep neural network to capture long-range inter-residue relationship and co-evolutionary information encoded in the sequence. Previous work[9, 28, 23, 20] has shown that with small-scale supervised training for downstream tasks, PLMs can capture some functional and structure properties of proteins, including secondary structure, binding residues[21], tertiary contact and protein structure[20, 40, 11]. Some work[29] also extends the PLMs to model a set of aligned sequences in a MSA using axial attention. AlphaFold2[15] also integrates the BERT-style objective to predict the masked elements of MSA sequence to improve MSA-based protein structure prediction.

### MSA-based Protein Structure Prediction

In the past few years, some methods[42, 41, 37] use coevolutionary features to predict protein structure using ResNet. Compared to directly using MSA as input, extracting coevolutionary features will lead to information loss. Recently MSA has been proposed as input directly using an encoder[24, 14] and decode the distance using 2D ResNet. Most methods use a ResNet-based model to infer inter-residue geometric constraints and build structure under those constraints. AlphaFold2[15] is the first successful end-to-end model supervised with structures. AlphaFold2 win the CASP14 and revolutionize protein structure prediction with so many new ideas, ranging from recycling its own prediction, over end-to-end training from MSA to structure, to clear ways of using triangular attention and a large database[38] with reliable accuracy estimates.

### Sequence-based Protein Structure Prediction

Since AlphaFold2 achieve remarkable performance at CASP14, some group are looking to speed up AlphaFold2 pipeline to reduce the computational expense, including model inference stage [2, 7] and homologs search stage[25, 13]. In nature a protein folds without knowledge of its sequence homologs, predicting protein structure based upon some non-natural conditions such as MSA or template does not reflect very well how a protein actually folds. In addition, not all proteins have enough sequence homologs, the MSA-based method does not perform well on those small-size protein families. Some groups[3, 39] devoted to study how to predict protein structure unaware of any sequence homologs. EvoGen[43] uses U-shaped neural network architecture to learn generalizable features across MSA and generate virtual MSA based on target sequence. OmegaFold[40] achieve fast structure prediction by combining the language model with structure module of AlphaFold2. ESMFold[20] presents ESM2 with 15 billion parameters as well as a better positional encoding to encode the inter-residue relationship and decode the coordinate using structure module similar to AlphaFold2.

### Protein complex prediction

In addition to accelerating AlphaFold2, many researchers have attempted to generalize AlphaFold2 to complex prediction. Some groups find that adding a gap or linker segment[25, 17, 5] between chains of complex can successfully model complex with pre-trained AlphaFold2, which outperforms some traditional template-based modeling and free docking[12, 6, 18]. DeepMind also presents AlphaFold-Multimer[10], a derived version of AlphaFold2 for multimer, has superior accuracy on complex structure prediction. However, compared to the powerful performance of AlphaFold2 on monomer, the accuracy of AlphaFold2 multimer on predicting the protein complex is far from satisfactory, especially for some antibodies.

### Antibody Structure Prediction

The modeling of CDR loops, especially the third CDR loop of the heavy chain (CDR H3) is a challenging topic. Inspired by the deep learning development in protein structure prediction, Some antibody-specific deep learning methods have been developed to improve CDR loop modeling accuracy. DeepAb[34] predicts a set of inter-residue geometric constraints and feeds the contraints to Rosetta to construct a complete antibody structure. ABlooper[1] is an end-to-end equivariant CDR loop prediction tool, the loop quality can be estimated under prediction. IgFold[31] extract sequence feature from AntiBERTy[32], update the sequence embedding and pairwise representation using triangle attention, and predict the backbone coordinates using IPA. IgFold achieves a fast and accurate antibody structure prediction.

## 3 Methods

As illustrated in Figure 1, the proposed antibody structure prediction pipeline mainly consists of four stages: 1) monomer structure prediction for each chain; 2) heavy-light chain feature fusion; 3) multimer structure prediction for the heavy-light chain complex; and 4) additional refinement with recycling iterations.

**Figure 1:**
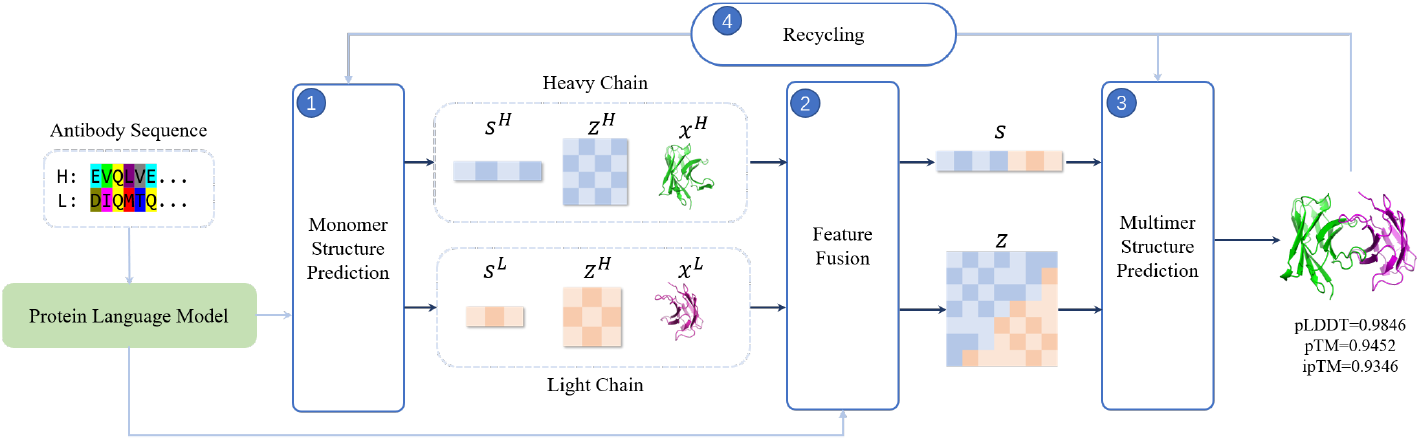
The overall workflow of antibody structure prediction pipeline.

### 3.1 Monomer Structure Prediction

It has been widely shown that pre-trained language models for protein sequences [27, 28, 30, 9] effectively capture the dependency among residues and thus provide meaningful feature embeddings. Such models are usually trained with a masked language modeling (MLM) loss, by feeding randomly masked amino-acid sequences into the network and forcing it to recover original ones via selfattention. Therefore, attention weights reflect how different residues interact with each other, which can be naturally used as initial pair features between residues.

Formally, for each heavy and light chain, we extract its final sequence embeddings and all-layer pairwise attention weights from the pre-trained ProtXLNet model [9]. Additionally, to explicitly distinguish heavy and light chains, we adopt two independent positional encoders with learnable embeddings, one per chain type. The monomer structure prediction for heavy and light chains consists of two stages: iterative updates of single and pair features, followed by 3D structure prediction. As illustrated in Figure 2, we mainly follow the architecture design in AlphaFold [15], with one minor difference in that the standard Evoformer stack (for updating MSA and pair features) is now replaced by a simplified Evoformer-Single stack for single sequence inputs (please refer to Appendix B for details). For efficiency, both heavy and light chains share the same set of model parameters for Evoformer-Single and structure modules, as we empirically find that this works well since: 1) heavy and light chains are similar in the overall conformation; and 2) independent learnable positional encodings provide chain-type specific information for monomer structure prediction.

**Figure 2:**
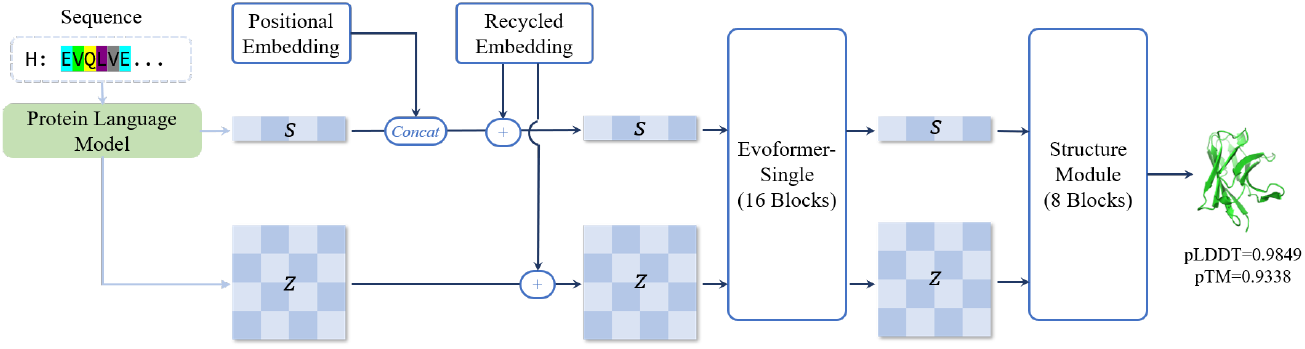
The network architecture for monomer and multimer structure prediction.

### 3.2 Heavy-Light Chain Feature Fusion

For multimer structure prediction of antibody heavy-light chain complex, we concatenate their aminoacid sequences without any delimiter. Correspondingly, each chain’s single features are concatenated along the sequence dimension as initial single features for multimer structure prediction. We re-use learnable positional encodings of heavy and light chains, so that the model is aware of each residue’s chain type and position.

As depicted in Figure 3, we divide pair features for multimer structure prediction into four parts, where diagonal blocks (*z^H^* and *z^L^*) are initialized by heavy and light chains’ pair features. For off-diagonal ones, we construct an extended antibody sequence where heavy and light chains are connected by 21 glycines as the linker. This sequence is then fed into the same ProtXLNet model, from which attention weights corresponding to inter-chain residue pairs are extracted to initialize pair features for multimer structure prediction.

**Figure 3:**
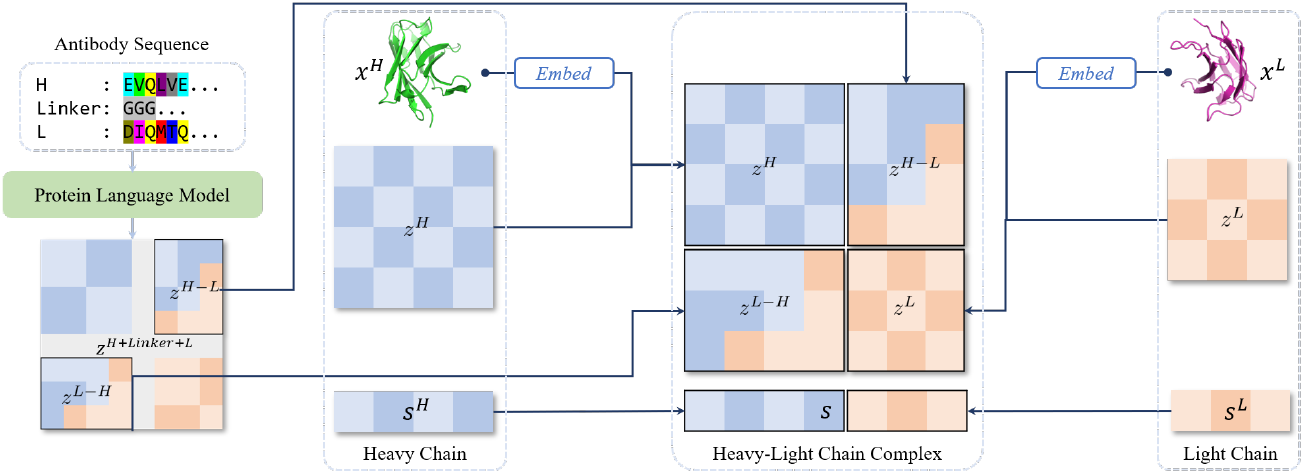
The fusion of heavy and light chain features for multimer structure prediction.

### 3.3 Multimer Structure Prediction

After heavy-light chain feature fusion, we feed initial single and pair features into another subnetwork consists of 16 Evoformer-Single blocks and 8 structure modules (with shared parameters) for multimer structure prediction. Despite the same network architecture, this sub-network do not share model parameters with that for monomer structure prediction, so that it concentrates on refining feature embeddings in the context of global conformation of heavy-light chain complex.

### 3.4 Recycling Iterations

So far, monomer structure prediction for heavy and light chains only consider its own amino-acid sequence, which ignores possible interaction between heavy and light chains. This may limit the iterative update of single and pair features, especially for residues within the interaction site, leading to sub-optimal initial features for multimer structure prediction. We resolve this issue by taking the predicted multimer structure for additional recycling iterations, as described in Algorithm 1. The detailed design of recycling embedding module can be found in Appendix C.

#### Algorithm 1 Model inference with recycling iterations

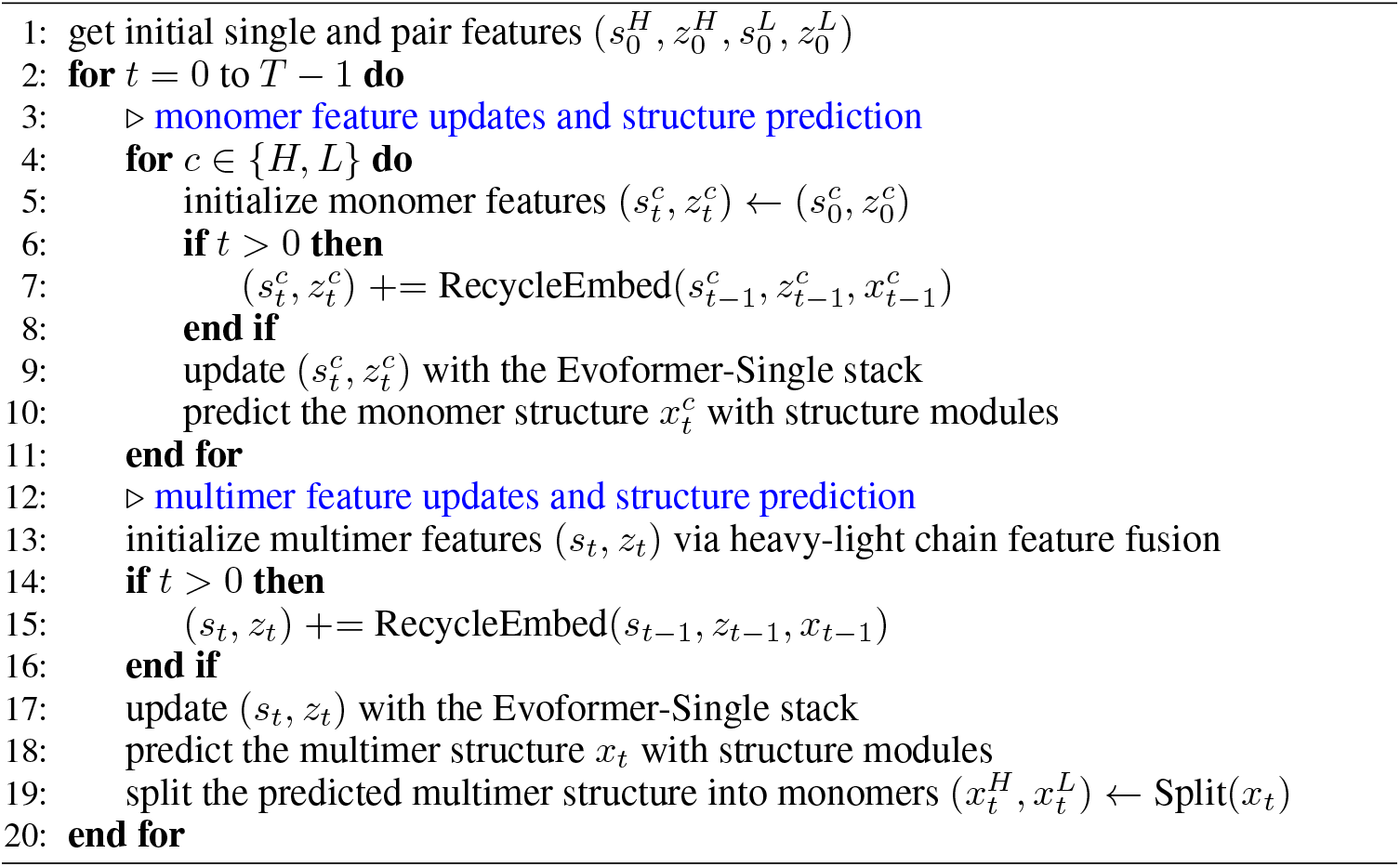

### 3.5 Model Optimization

As mentioned earlier, the proposed model can be trained with both single-chain and double-chain antibody structures. This is achieved by adopting a mixture of monomer and multimer structure prediction loss functions, given by:

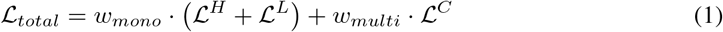

where 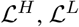, and 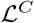 are loss functions for heavy chain, light chain, and heavy-light chain complex, respectively. We introduce *w_mono_* and *w_multi_* to balance these loss functions: if both chains exist, then *w_mono_* = 0.25 and *w_multi_* = 0.5; otherwise, we have *w_mono_* = 1 and *w_multi_* = 0. This ensures that both heavy and light chains contribute equally in the loss computation. Please refer to Appendix D for additional details on above loss functions.

## 4 Results

### Datasets

We construct train/valid/test subsets from the SAbDab database [36] in a temporally separated manner, similar to IgFold [31]. Specifically, we gather all the experimentally determined structures released before 2021/03/31 as training samples, which include 7591 heavy-light chain complexes, 1440 heavy-chain only, and 507 light-chain only samples. We conduct hyper-parameter tuning and model selection on the validation subset, consisting of structures released between 2021/04/01 and 2021/06/30. This guarantees a fair comparison on the IgFold-Ab benchmark (67 antibodies) since there is no temporal overlap. In addition, we build two larger non-redundant benchmarks (sequence identity lower than 95%) containing 235 antibodies and 69 nanobodies released during the first 6 months in 2022, namely SAbDab-22H1-Ab/Nano. We do not remove redundant structures with identical sequences from the training subset, since it is beneficial to expose the model with alternative conformations during training [1].

### Training

We adopt the Adam optimizer with a fixed learning rate of 3e-4 and set the batch size to 32. We firstly train the model without recycling iterations for 50 epochs, and then fine-tune it for another 100 epochs with the number of recycling iterations (T) set to 2. We maintain the exponential moving average of model parameters with *α* = 0.999 and use this model for evaluation. The optimal model is selected based on full-atom RMSD scores on the validation subset.

### Evaluation

Baseline methods include antibody-specific structure prediction (ABodyBuilder [19], DeepAb [34], ABlooper [1], NanoNet [8] and IgFold [31]) and general protein structure prediction, either MSA-based (AlphaFold [15] and AlphaFold-Multimer [10]) or MSA-free (HelixFold-Single [11], ESMFold [20], and OmegaFold [40]).

For antibody structure prediction, we report backbone RMSD in different framework and CDR regions on IgFold-Ab (Table 1) and SAbDab-22H1-Ab (Table 2) benchmarks. The proposed method achieves the lowest RMSD in almost all regions, with the only exception of CDR-L2. Notably, the RMSD of most flexible CDR-H3 is reduced from 2.99Å to 2.74Å on IgFold-Ab, and from 3.18Å to 3.03Å on SAbDab-22H1-Ab, indicating that our method accurately predict the overall conformation of this region. Besides, we compare OCD (orientational coordinate distance) [22] to verify how well the relative position between heavy and light chains is estimated. Again, our method consistently outperforms all the baselines on both benchmarks.

**Table 1:** Antibody structure prediction accuracy on the IgFold-Ab benchmark.

| Method | OCD | H-Fr | H1<sup>3</sup> | H2 | H3 | L-Fr | L1 | L2 | L3 |
| --- | --- | --- | --- | --- | --- | --- | --- | --- | --- |
| ABodyBuilder [19] | 4.90 | 0.54 | 1.10 | 0.94 | 3.75 | 0.43 | 1.07 | 0.58 | 1.37 |
| ABlooper [1] | 4.53 | 0.51 | 0.95 | 0.82 | 3.20 | 0.45 | 0.99 | 0.59 | 1.15 |
| DeepAb [34] | 3.60 | 0.43 | 0.80 | 0.74 | 3.28 | 0.38 | 0.86 | 0.45 | 1.11 |
| IgFold [31] | 3.77 | 0.45 | 0.80 | 0.75 | 2.99 | 0.45 | 0.83 | 0.51 | 1.07 |
| AlphaFold-Multimer [10] | 3.69 | 0.43 | 0.75 | 0.69 | 3.02 | 0.39 | 0.82 | **0.41** | 1.13 |
| HelixFold-Single [11] | - | 0.56 | 0.85 | 0.95 | 5.01 | 0.51 | 1.10 | 0.57 | 1.60 |
| ESMFold [20] | - | 0.51 | 0.84 | 0.91 | 4.10 | 0.43 | 1.16 | 0.52 | 1.44 |
| OmegaFold [40] | - | 0.47 | 0.75 | 0.74 | 3.70 | 0.41 | 0.93 | 0.43 | 1.35 |
| tFold-Ab | **3.21** | **0.41** | **0.69** | **0.63** | **2.74** | **0.37** | **0.75** | 0.43 | **0.92** |

**Table 2:** Antibody structure prediction accuracy on the SAbDab-22H1-Ab benchmark.

| Method | OCD | H-Fr | H1 | H2 | H3 | L-Fr | L1 | L2 | L3 |
| --- | --- | --- | --- | --- | --- | --- | --- | --- | --- |
| DeepAb [34] | 5.93 | 0.58 | 1.07 | 0.90 | 3.79 | 0.61 | 0.93 | 0.73 | 1.41 |
| IgFold [31] | 5.66 | 0.61 | 1.06 | 0.93 | 3.46 | 0.60 | 0.90 | 0.71 | 1.31 |
| AlphaFold-Multimer [10] | 5.32 | 0.57 | 1.02 | **0.85** | 3.18 | 0.57 | 0.84 | 0.69 | 1.23 |
| HelixFold-Single [11] | - | 0.69 | 1.16 | 1.12 | 5.70 | 0.66 | 1.09 | 0.80 | 1.79 |
| ESMFold [20] | - | 0.62 | 1.11 | 1.04 | 4.82 | 0.60 | 1.12 | 0.73 | 1.71 |
| OmegaFold [40] | - | 0.61 | 1.04 | 0.87 | 4.35 | 0.57 | 0.91 | **0.68** | 1.43 |
| tFold-Ab | **5.10** | **0.55** | **0.97** | 0.86 | **3.03** | **0.54** | **0.81** | 0.71 | **1.19** |

Furthermore, we report DockQ [4] to measure the docking pose of heavy and light chains in Table 3. Although our method achieves better backbone prediction quality (LRMS and iRMS), its Fnat score (involves both backbone and side-chain atoms) is inferior to AlphaFold-Multimer, leading to the lower DockQ score. This indicates that the prediction of side-chain conformation has yet to improve.

**Table 3:**
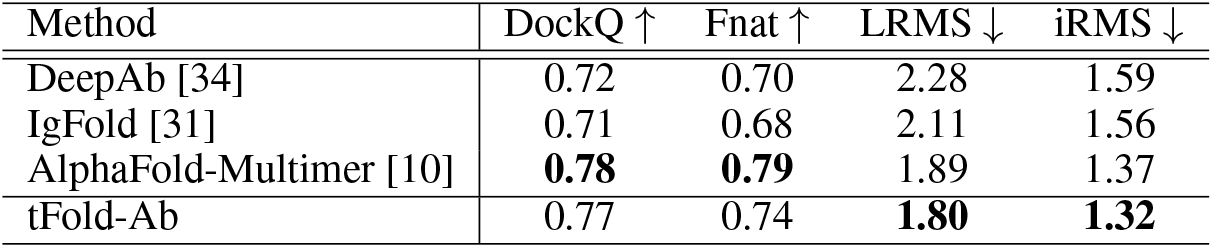
DockQ evaluation results on the SAbDab-22H1-Ab benchmark.

| Method | DockQ ↑ | Fnat ↑ | LRMS ↓ | iRMS ↓ |
| --- | --- | --- | --- | --- |
| DeepAb [34] | 0.72 | 0.70 | 2.28 | 1.59 |
| IgFold [31] | 0.71 | 0.68 | 2.11 | 1.56 |
| AlphaFold-Multimer [10] | **0.78** | **0.79** | 1.89 | 1.37 |
| tFold-Ab | 0.77 | 0.74 | **1.80** | **1.32** |

**Table 4:**
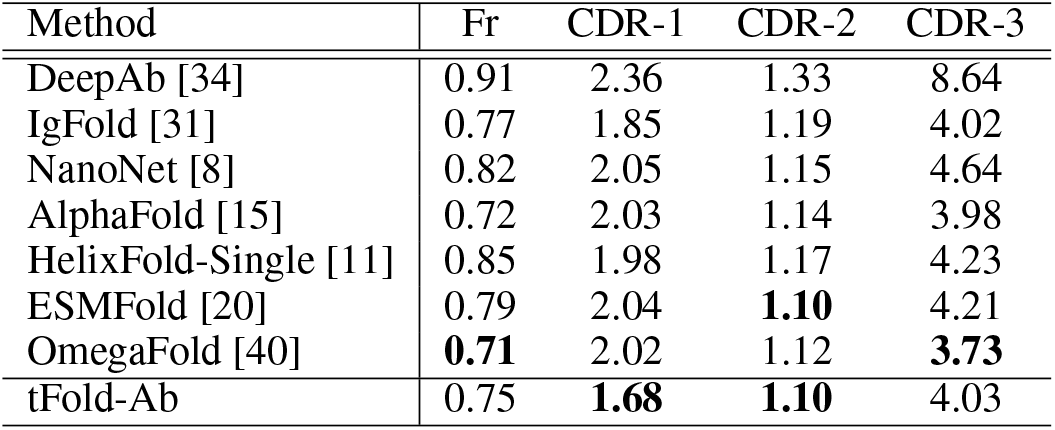
Nanobody structure prediction accuracy on the SAbDab-22H1-Nano benchmark.

For nanobody structure prediction, we omit the IgFold-Nano benchmark due to limited number of test samples (21 nanobodies), and only report performance on SAbDab-22H1-Nano. Our method achieves the lowest RMSD in both CDR-1 and CDR-2 regions, but does not work well in framework and CDR-3 regions. This may be caused by the relatively low fraction of nanobodies (< 20%) in the training subset, and should be further investigated in future works.

## 5 Discussion and Limitations

In this paper, we propose a fully end-to-end architecture for antibody and nanobody structure prediction, which achieves state-of-the-art performance in various benchmarks. Despite its promising preliminary results, there are still much to improve in the current model. The choice of pre-trained language models is not well studied; recent large-scale pre-trained language models, *e.g*., ESM-2 [20], may further improve the antibody structure prediction accuracy, and should be investigated in future works. Furthermore, the OAS database [26] contains large-scale antibody sequences without available experimentally determined structures, and may be exploited through self-distillation training, but the performance improvement remains uncertain. Finally, the interaction between antibody and antigen is not explicitly formulated by the current model; however, such interaction is vital in determining the stable conformation of CDR loops. It would be interesting to develop a unified model for modelling such antibody-antigen binding mechanism, thus enabling the rational design of antibodies.

## A Data Availability

All the FASTA sequences, native structures, and tFold-Ab predicted structures for three benchmarks (IgFold-Ab and SAbDab-22H1-Ab/Nano) used in this paper can be downloaded from https://drive.google.com/file/d/15C5hbd0mGgOcdXXb0x5Af2COVy7nXzpt/view?usp=sharing.

## B Evoformer-Single Stack

As mentioned earlier, our method only takes the amino-acid sequence itself as inputs, without any sequence homologs. Therefore, for iterative updates of single and pair features, we simplify AlphaFold’s Evoformer stack [15] (originally designed for updating MSA and pair features) to handle single sequence inputs, as illustrated in Algorithm 2.

### Algorithm 2 Evoformer-Single Stack

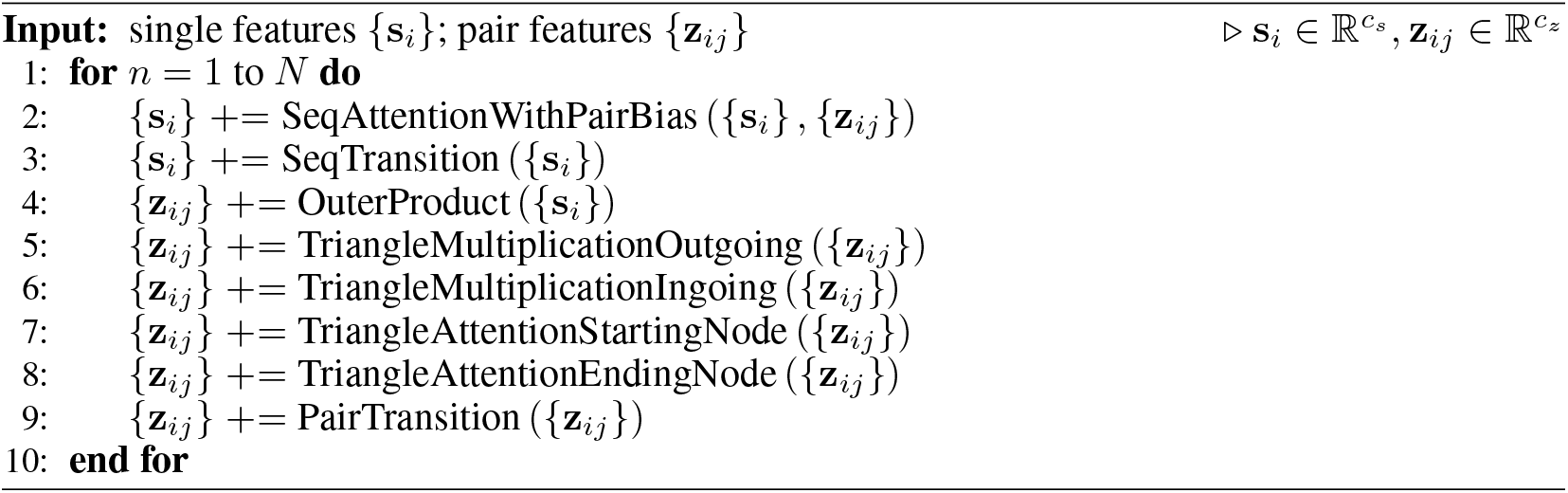

Specifically, “MSAColumnAttention” in the original Evoformer block is removed since calculating attention weights among sequence homologs is no longer applicable for single sequence inputs. Other MSA-related modules are simplified correspondingly, while modules only involving pair features remain unchanged.

In Algorithm 3, single features are updated via the gated self-attention mechanism along the sequence dimension to formulate the inter-residue interaction. Additionally, pair features are linearly projected into bias terms, which are then imposed into attention weights. This allows updating single features under the guidance of inter-residue pair features.

### Algorithm 3 SeqAttentionWithPairBias

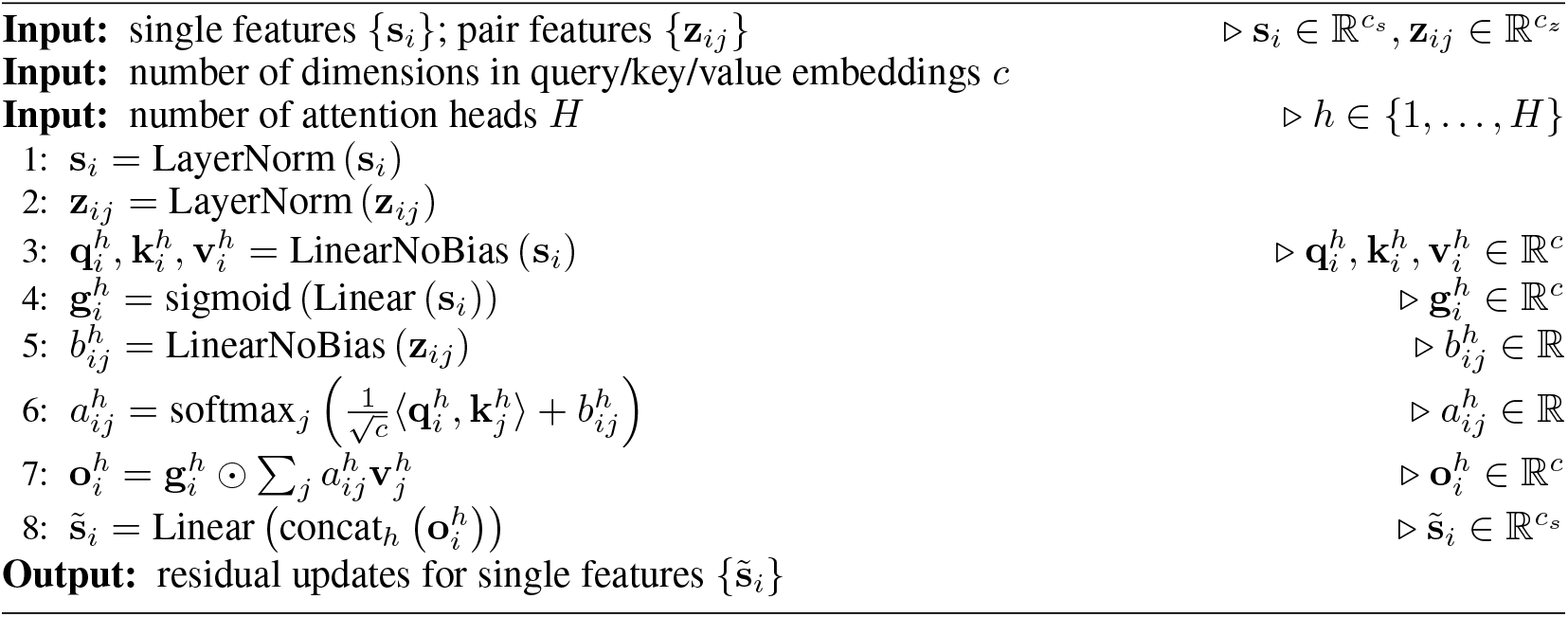

In Algorithm 4, single features are updated via a two-layer feed-forward network. We set the number of dimensions in hidden embeddings as *c* = 4*c_s_*, same as AlphaFold. Afterwards, pair features are updated based on the outer-product of single features, as described in Algorithm 5.

### Algorithm 4 SeqTransition

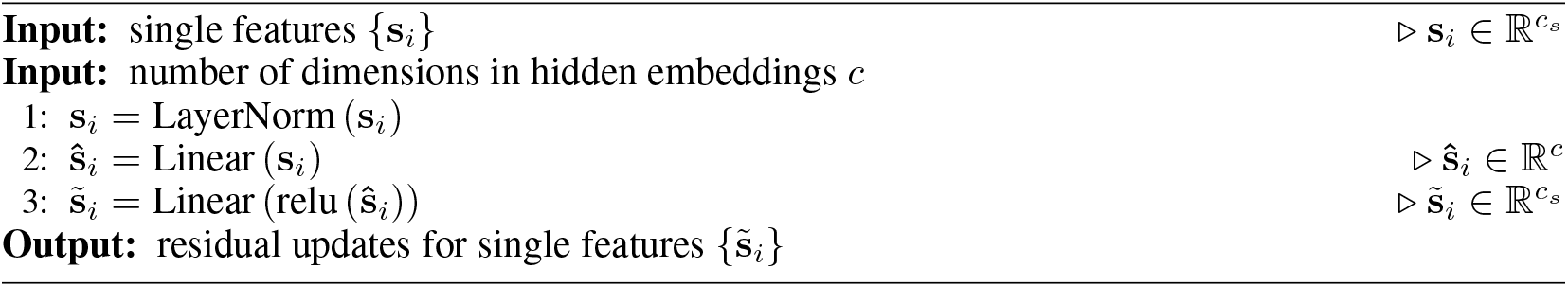

### Algorithm 5 OuterProduct

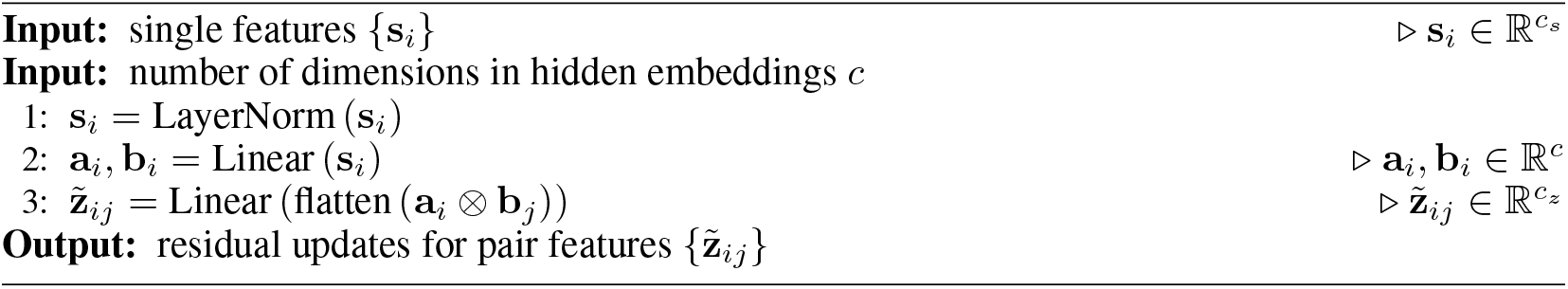

## C Recycling Embedding Module

In order to utilize feature embeddings and structure predictions from the previous iteration (as in Algorithm 1), we slightly modify the original “RecyclingEmbedder” module in AlphaFold [15] for single sequence inputs. The modified recycling embeddings module is as described in Algorithm 6: Specifically, single and pair features from the previous iteration are normalized to produce residue update terms for the current iteration. Atomic coordinates of *C_β_* (*C_α_* for glycines) atoms are extracted from the predicted structure, from which pairwise Euclidean distance is computed. Such distance is then discretized into histogram bins to generate one-hot encodings for the final linear projection.

### Algorithm 6 RecycleEmbed

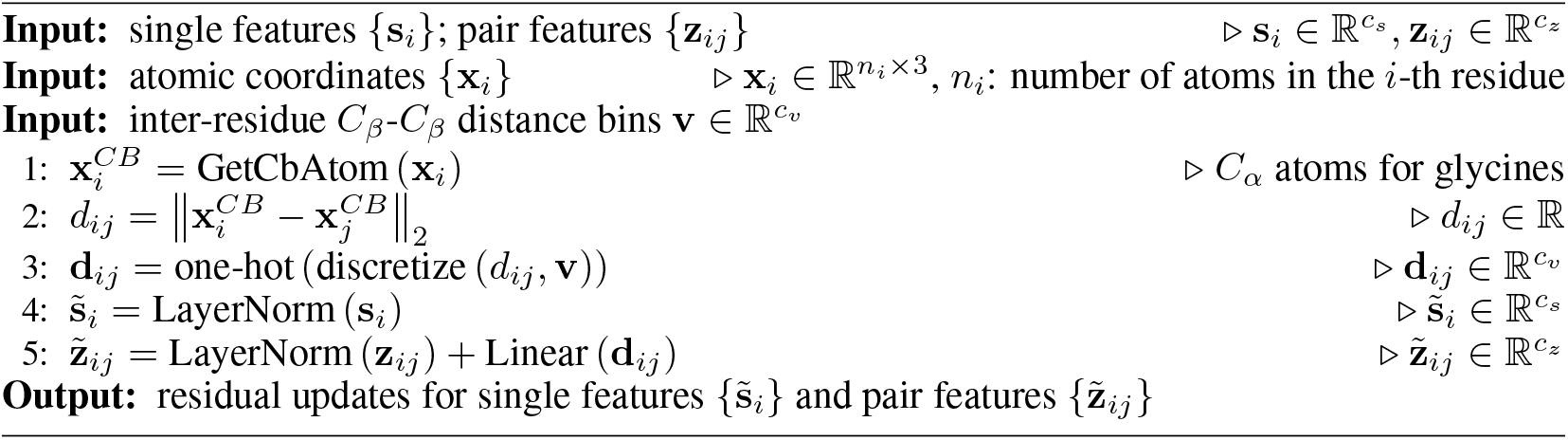

## D Loss Function

As described in Section 3.5, the overall loss function consists of a mixture of monomer and multimer losses, denoted as 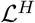 (heavy chain), 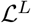 (light chain), and 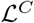 (heavy-light chain complex). Each loss term is defined on the predicted monomer/multimer structure and auxiliary predictions (*e.g*., inter-residue distance), sharing the same form:

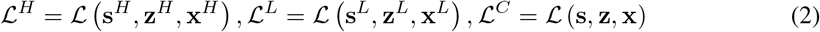

Therefore, we take the multimer loss function 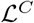 as an example, and describe its detailed loss terms. Concretely, this loss function constitutes of following terms:

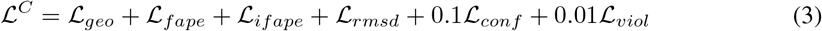

### Inter-residue geometric loss 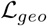

To provide more direct supervision in the Evoformer-Single stack, we add four auxiliary heads (implemented as feed-forward layers) on the top of final pair features for predicting inter-residue distance and angles, as defined in trRosetta [42]. This includes:

- 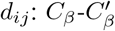 distance
- 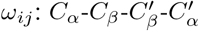 dihedral angle
- 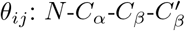 dihedral angle
- 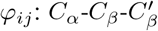 planar angle

Each prediction head outputs probabilistic estimations of above distance and angles, denoted as 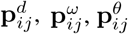, and 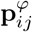. We then calculate the cross-entropy loss for each term and sum them up to as the final inter-residue geometric loss:

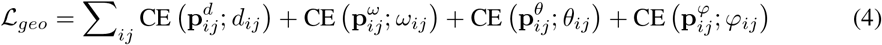

### Frame aligned point error 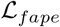

This is identical to the FAPE loss used in AlphaFold [15]. After reconstructing full-atom 3D coordinates from per-residue backbone frame and torsion angle predictions, each atom is projected into all the local frames (both backbone and side-chain) in the ground-truth and predicted structures for comparison.

### Interface frame aligned point error 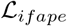

This is identical to the second part of FAPE loss used in AlphaFold-Multimer [10], which is applied to inter-chain residue pairs and clamped at 30Å. Please note that this loss is only computed over multimer structure predictions.

### Coordinate RMSD loss 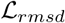

In order to better estimate the overall conformation, we calculate the coordinate RMSD (root-of-mean-squared-deviation) loss between ground-truth and predicted structures after alignment. The optimal alignment is determined by the Kabsch algorithm [16] for finding the optimal rotation and translation between two sets of point clouds. For the *i*-th residue’s *j*-th atom, we denote its 3D coordinate in the ground-truth structure as 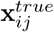, and 3D coordinate in the aligned predicted structures as 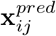. The coordinate RMSD loss is defined as:

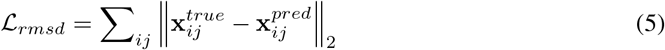

### Confidence loss 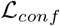

This includes loss functions for pLDDT and pTM predictions, same as AlphaFold [15]. We detach single and pair features before estimating pLDDT and pTM scores, similar to IgFold [31], to prevent the model from generating problematic structures whose lDDT and TM-score can be accurately predicted.

### Structure violation loss 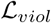

Similar to AlphaFold [15], we introduce penalty terms for incorrect peptide bond length and angles, as well as steric clashes between non-bonded atoms. For multimer structure prediction, we do not penalize the bond length and angle between the last residue in the heavy chain and first residue in the light chain, since there is no peptide bond between them. Besides, we normalize the steric clash loss by the number of non-bonded atom pairs in clash to stablize the model optimization, as suggested in AlphaFold-Multimer [10].

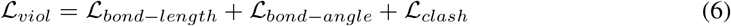

## E Implementation Details

We set the number of dimensions in single and pair features as *c_s_* = 384 and *c_z_* = 256. The network consists of two Evoformer-Single & structure module blocks for monomer and multimer structure prediction, respectively. Each block contains 16 Evorformer-Single blocks (without shared parameters) and 8 structure modules (with shared parameters). To ease the optimization difficulty, we start with 2 structure modules and gradually increase the depth until maximum during the training process (one more structure module every two epochs). The model is trained with 32 A100 GPUs for a total of 150 epochs, which takes roughly 80 hours to finish. The first 50 epochs are trained without the structure violation loss; it is only enabled in the second stage of model training. At the end of each epoch, we evaluate the model on the validation set and record its full-atom RMSD. Once finished, the checkpoint with lowest validation full-atom RMSD is selected as the final one. The validation full-atom RMSD also serves as the criterion for hyper-parameter tuning, *e.g*., learning rate and batch size.

## F Inference Speed

One major advantage of our proposed method is that the time-consuming MSA search procedure is no longer needed, due to the utilization of pre-trained language models. In addition, our model formulates both backbone and side-chain conformations with a unified neural network, while previous antibody structure prediction methods, *e.g*., DeepAb [34] and IgFold [31], rely on Rosetta-based energy minimization to predict side-chain structures. AlphaFold-Multimer [10] predicts full-atom structures with a single forward pass, but the computational complexity of its Evoformer stack is much larger than ours.

In Figure 4, we report the time consumption of various antibody structure prediction methods on the SAbDab-22H1-Ab benchmark. All the run time is measured on a single A100 GPU with 21 CPU cores. For AlphaFold-Multimer, all the sequence and template databases are stored on a distributed file system (Ceph), thus the time consumption may be further reduced if local SSD disks are used instead. Therefore, we also report its execution time with MSA and template search procedures excluded, denoted as “AF-Multimer (NN-only)”. Our proposed method is able to predict full-atom antibody structures within 3 seconds, only slower than “IgFold (w/o PyRosetta)” which only produces backbone structure predictions.

**Figure 4:**
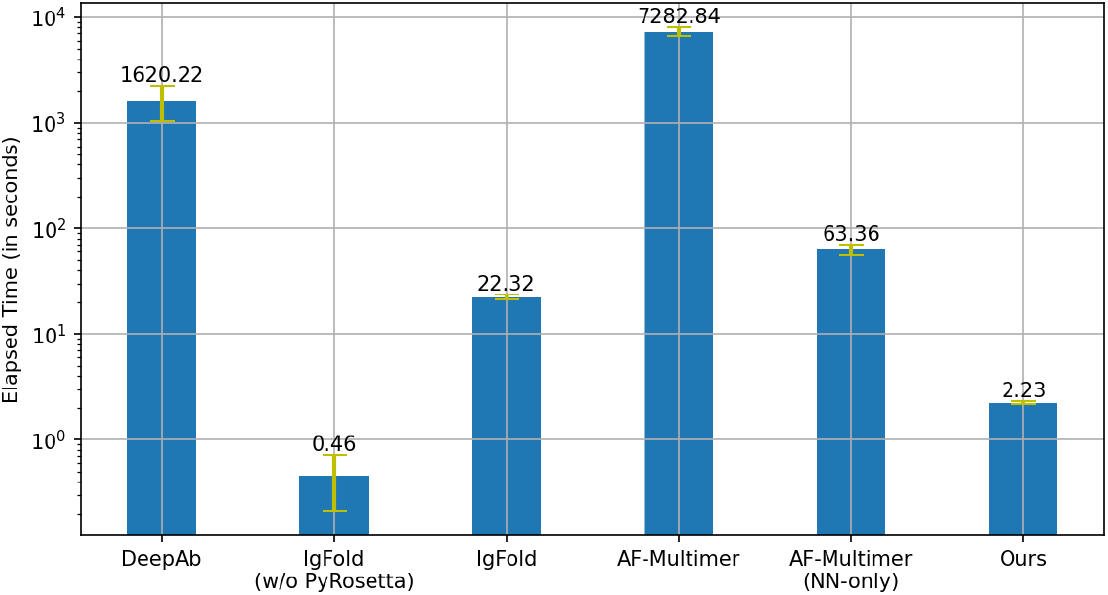
Comparison on the antibody structure prediction time of various methods on the SAbDab-22H1-Ab benchmark. By default, IgFold uses PyRosetta to generate side-chain conformations from backbone structure predictions. AlphaFold-Multimer runs all the five candidate models and then selects the optimal one based on the combination of pTM and ipTM scores. We report the execution time excluding MSA and template search of AlphaFold-Multimer as “AF-Multimer (NN-only)”.

## Footnotes

3 We use “H1” as the abbreviation for “CDR-H1” to save space.

## Notes

https://drive.google.com/file/d/15C5hbd0mGgOcdXXb0x5Af2COVy7nXzpt/view?usp=sharing

### Competing Interest Statement

The authors have declared no competing interest.

## References

[1] B. Abanades, G. Georges, A. Bujotzek, and C. M. Deane. Ablooper: Fast accurate antibody cdr loop structure prediction with accuracy estimation. Bioinformatics, 38(7):1877–1880, 2022.

[2] G. Ahdritz, N. Bouatta, S. Kadyan, Q. Xia, W. Gerecke, and M. AlQuraishi. OpenFold, 11 2021.

[3] M. AlQuraishi. End-to-end differentiable learning of protein structure. Cell systems, 8(4):292–301, 2019.

[4] S. Basu and B. Wallner. Dockq: A quality measure for protein-protein docking models. PLOS ONE, 11(8):1–9, 08 2016.

[5] P. Bryant, G. Pozzati, and A. Elofsson. Improved prediction of protein-protein interactions using alphafold2. Nature communications, 13(1):1–11, 2022.

[6] R. Chen, L. Li, and Z. Weng. Zdock: an initial-stage protein-docking algorithm. Proteins: Structure, Function, and Bioinformatics, 52(1):80–87, 2003.

[7] S. Cheng, R. Wu, Z. Yu, B. Li, X. Zhang, J. Peng, and Y. You. Fastfold: Reducing alphafold training time from 11 days to 67 hours, 2022.

[8] T. Cohen, M. Halfon, and D. Schneidman-Duhovny. NanoNet: Rapid and accurate end-to-end nanobody modeling by deep learning. Front. Immunol., 13:958584, Aug. 2022.

[9] A. Elnaggar, M. Heinzinger, C. Dallago, G. Rehawi, Y. Wang, L. Jones, T. Gibbs, T. Feher, C. Angerer, M. Steinegger, et al. Prottrans: towards cracking the language of lifes code through self-supervised deep learning and high performance computing. IEEE transactions on pattern analysis and machine intelligence, 2021.

[10] R. Evans, M. O’Neill, A. Pritzel, N. Antropova, A. W. Senior, T. Green, A. Žídek, R. Bates, S. Blackwell, J. Yim, et al. Protein complex prediction with alphafold-multimer. BioRxiv, 2021.

[11] X. Fang, F. Wang, L. Liu, J. He, D. Lin, Y. Xiang, X. Zhang, H. Wu, H. Li, and L. Song. Helixfold-single: Msa-free protein structure prediction by using protein language model as an alternative. arXiv preprint arXiv:2207.13921, 2022.

[12] A. Guerler, B. Govindarajoo, and Y. Zhang. Mapping monomeric threading to protein–protein structure prediction. Journal of chemical information and modeling, 53(3):717–725, 2013.

[13] L. Hong, S. Sun, L. Zheng, Q. Tan, and Y. Li. fastmsa: Accelerating multiple sequence alignment with dense retrieval on protein language. bioRxiv, 2021.

[14] F. Ju, J. Zhu, B. Shao, L. Kong, T.-Y. Liu, W.-M. Zheng, and D. Bu. Copulanet: Learning residue co-evolution directly from multiple sequence alignment for protein structure prediction. Nature communications, 12(1):1–9, 2021.

[15] J. Jumper, R. Evans, A. Pritzel, T. Green, M. Figurnov, O. Ronneberger, K. Tunyasuvunakool, R. Bates, A. Žídek, A. Potapenko, A. Bridgland, C. Meyer, S. A. A. Kohl, A. J. Ballard, A. Cowie, B. Romera-Paredes, S. Nikolov, R. Jain, J. Adler, T. Back, S. Petersen, D. Reiman, E. Clancy, M. Zielinski, M. Steinegger, M. Pacholska, T. Berghammer, S. Bodenstein, D. Silver, O. Vinyals, A. W. Senior, K. Kavukcuoglu, P. Kohli, and D. Hassabis. Highly accurate protein structure prediction with alphafold. Nature, 596(7873):583–589, Aug 2021.

[16] W. Kabsch. A solution for the best rotation to relate two sets of vectors. Acta Crystallographica Section A: Crystal Physics, Diffraction, Theoretical and General Crystallography, 32(5):922–923, 1976.

[17] J. Ko and J. Lee. Can alphafold2 predict protein-peptide complex structures accurately? BioRxiv, 2021.

[18] D. Kozakov, D. R. Hall, B. Xia, K. A. Porter, D. Padhorny, C. Yueh, D. Beglov, and S. Vajda. The cluspro web server for protein–protein docking. Nature protocols, 12(2):255–278, 2017.

[19] J. Leem, J. Dunbar, G. Georges, J. Shi, and C. M. Deane. ABodyBuilder: Automated antibody structure prediction with data-driven accuracy estimation. MAbs, 8(7):1259–1268, Oct. 2016.

[20] Z. Lin, H. Akin, R. Rao, B. Hie, Z. Zhu, W. Lu, A. dos Santos Costa, M. Fazel-Zarandi, T. Sercu, S. Candido, et al. Language models of protein sequences at the scale of evolution enable accurate structure prediction. bioRxiv, 2022.

[21] M. Littmann, M. Heinzinger, C. Dallago, K. Weissenow, and B. Rost. Protein embeddings and deep learning predict binding residues for various ligand classes. Scientific Reports, 11(1):1–15, 2021.

[22] N. A. Marze, S. Lyskov, and J. J. Gray. Improved prediction of antibody VL-VH orientation. Protein Eng. Des. Sel., 29(10):409–418, Oct. 2016.

[23] J. Meier, R. Rao, R. Verkuil, J. Liu, T. Sercu, and A. Rives. Language models enable zero-shot prediction of the effects of mutations on protein function. Advances in Neural Information Processing Systems, 34:29287–29303, 2021.

[24] C. Mirabello and B. Wallner. Rawmsa: End-to-end deep learning using raw multiple sequence alignments. PloS one, 14(8):e0220182, 2019.

[25] M. Mirdita, K. Schütze, Y. Moriwaki, L. Heo, S. Ovchinnikov, and M. Steinegger. Colabfold: making protein folding accessible to all. Nature Methods, pages 1–4, 2022.

[26] T. H. Olsen, F. Boyles, and C. M. Deane. Observed antibody space: A diverse database of cleaned, annotated, and translated unpaired and paired antibody sequences. Protein Science, 31(1):141–146, 2022.

[27] R. Rao, N. Bhattacharya, N. Thomas, Y. Duan, P. Chen, J. Canny, P. Abbeel, and Y. Song. Evaluating protein transfer learning with tape. Advances in neural information processing systems, 32, 2019.

[28] R. Rao, J. Meier, T. Sercu, S. Ovchinnikov, and A. Rives. Transformer protein language models are unsupervised structure learners. Biorxiv, 2020.

[29] R. M. Rao, J. Liu, R. Verkuil, J. Meier, J. Canny, P. Abbeel, T. Sercu, and A. Rives. Msa transformer. In International Conference on Machine Learning, pages 8844–8856. PMLR, 2021.

[30] A. Rives, J. Meier, T. Sercu, S. Goyal, Z. Lin, J. Liu, D. Guo, M. Ott, C. L. Zitnick, J. Ma, et al. Biological structure and function emerge from scaling unsupervised learning to 250 million protein sequences. Proceedings of the National Academy of Sciences, 118(15):e2016239118, 2021.

[31] J. A. Ruffolo and J. J. Gray. Fast, accurate antibody structure prediction from deep learning on massive set of natural antibodies. Biophysical Journal, 121(3):155a–156a, 2022.

[32] J. A. Ruffolo, J. J. Gray, and J. Sulam. Deciphering antibody affinity maturation with language models and weakly supervised learning. arXiv preprint arXiv:2112.07782, 2021.

[33] J. A. Ruffolo, C. Guerra, S. P. Mahajan, J. Sulam, and J. J. Gray. Geometric potentials from deep learning improve prediction of cdr h3 loop structures. bioRxiv, 2020.

[34] J. A. Ruffolo, J. Sulam, and J. J. Gray. Antibody structure prediction using interpretable deep learning. Patterns, 3(2):100406, 2022.

[35] V. G. Satorras, E. Hoogeboom, and M. Welling. E(n) equivariant graph neural networks. In M. Meila and T. Zhang, editors, Proceedings of the 38th International Conference on Machine Learning, volume 139 of Proceedings of Machine Learning Research, pages 9323–9332. PMLR, 18–24 Jul 2021.

[36] C. Schneider, M. I. J. Raybould, and C. M. Deane. SAbDab in the age of biotherapeutics: updates including SAbDab-nano, the nanobody structure tracker. Nucleic Acids Research, 50(D1):D1368–D1372, 11 2021.

[37] A. W. Senior, R. Evans, J. Jumper, J. Kirkpatrick, L. Sifre, T. Green, C. Qin, A. Žídek, A. W. Nelson, A. Bridgland, et al. Improved protein structure prediction using potentials from deep learning. Nature, 577(7792):706–710, 2020.

[38] M. Varadi, S. Anyango, M. Deshpande, S. Nair, C. Natassia, G. Yordanova, D. Yuan, O. Stroe, G. Wood, A. Laydon, et al. Alphafold protein structure database: massively expanding the structural coverage of protein-sequence space with high-accuracy models. Nucleic acids research, 50(D1):D439–D444, 2022.

[39] W. Wang, Z. Peng, and J. Yang. Single-sequence protein structure prediction using supervised transformer protein language models. bioRxiv, 2022.

[40] R. Wu, F. Ding, R. Wang, R. Shen, X. Zhang, S. Luo, C. Su, Z. Wu, Q. Xie, B. Berger, et al. High-resolution de novo structure prediction from primary sequence. BioRxiv, 2022.

[41] J. Xu. Distance-based protein folding powered by deep learning. Proceedings of the National Academy of Sciences, 116(34):16856–16865, 2019.

[42] J. Yang, I. Anishchenko, H. Park, Z. Peng, S. Ovchinnikov, and D. Baker. Improved protein structure prediction using predicted interresidue orientations. Proceedings of the National Academy of Sciences, 117(3):1496–1503, 2020.

[43] J. Zhang, S. Liu, M. Chen, H. Chu, M. Wang, Z. Wang, J. Yu, N. Ni, F. Yu, D. Chen, et al. Few-shot learning of accurate folding landscape for protein structure prediction. arXiv preprint arXiv:2208.09652, 2022.

